# Equine Parentage Testing with SNPs and beyond: developing a platform independent panel

**DOI:** 10.64898/2025.12.16.694674

**Authors:** Rajesh Patidar, Ghadeer El-Ghabeish, Mais Abdel Wahhab, Heba Aqeel, Omar Adas Blanco, Mustafa K Khokha, Saquib A Lakhani

## Abstract

Centuries of selective equine breeding have led to a remarkable diversity of horse breeds each with unique traits. To maintain these breeds, studbooks require parentage testing to guarantee lineage. To determine parentage, the industry assays length polymorphisms in short tandem repeats (STRs) established by the International Society of Animal Genetics (ISAG). However, STRs have significant limitations including challenges with accurately determining genotype, relatively low number of STRs assayed, and low throughput. For these reasons, most other animal breeds including cattle, dog, goats, and sheep use single nucleotide polymorphisms (SNPs) for parentage identification. To this end, the ISAG has introduced a panel of SNPs. Here, we further demonstrate the utility of equine SNP genotyping. Importantly, we show that a number of the ISAG SNPs are not compatible with widely-used, short-read, next generation sequencing (NGS) technologies. Fortunately, SNPs are abundant, and we propose an alternative panel that is platform independent and are all ISAG proposed SNPs. We demonstrate that SNP parentage genotyping is extremely robust and enables additional assays including breed identification and can be readily expanded to also assay equine genetic diseases.

## INTRODUCTION

The global horse trade is a multibillion-dollar industry with numerous breeds exhibiting a plethora of desirable traits. Central to this are breed associations and studbooks, with the Thoroughbred studbook dating back to 1791 as the oldest recorded example (Weatherbys, 2025). Registration within these organizations is essential for upholding pedigree integrity and breed purity, ensuring that buyers, breeders, and owners can trust the lineage of their horses.

However, traditional registration systems rely on a degree of trust. Errors or discrepancies may occur, whether unintentionally, such as when mares are exposed to multiple stallions in shared environments, or deliberately, to manipulate pedigree records for perceived performance or commercial advantages. Additionally, the introduction of bloodlines from other breeds, sometimes intended to enhance specific traits, can further complicate the maintenance of genetic integrity. To address these challenges and reinforce the reliability of pedigree documentation, many horse breed associations now require genetic testing to verify parentage.

In 1995, the first STR (short tandem repeats) markers were published for thoroughbred parentage testing (Binns, 1995). STRs are generally di-nucleotide repeats (although they can have repeats of 1-6 base pairs) that are highly polymorphic because of slippage of the DNA polymerase during replication.

Additionally, they can be easily assayed by PCR and measurement of amplicon size via electrophoresis. After the first STRs were defined, an analysis of 4,803 Quarter Horses showed that microsatellites achieved 98.2% accuracy for parentage verification (Bowling, 1997). This development led to the International Society of Animal Genetics (ISAG) establishing 9 STR loci for equine parentage verification. This was subsequently increased to 12 STR loci in 2011, which remains the ISAG standard (van de Goor, 2011; Bellone, 2020).

However, there are numerous disadvantages to using STR markers compared to contemporary technologies. Due to errors in PCR amplification of STRs, the stutter peaks complicate size interpretation and can lead to false ascertainment of genotype as well as discordance between laboratories. In addition, allelic dropout has been reported leading to assignment of false homozygosity. Finally, STRs can be challenging to multiplex and automate scoring (Ishige, 2023). The ISAG recorded that only 72% of laboratories answered the parentage exclusion question correctly in 2017 (ISAG, 2017). There was little improvement in 2021; the ISAG stated that only 76% of laboratories entering the ISAG parentage test correctly assigned parentage (ISAG, 2021).

In contrast, SNP testing for parentage confers a number of advantages. First and foremost, they are highly accurate as hundreds to thousands to millions of loci can be assayed, such that parentage can be definitively determined even if a small percentage fail (i.e. SNP parentage determination is highly certain). Second, they can be highly multiplexed and automatically scored allowing for standardization (Holl, 2017). Third, hundreds of samples can be processed simultaneously, over a short time span, with same day results (Holl, 2017). Fourth, SNPs are more robust compared to STRs and can effectively assayed even when DNA extraction is sub-optimal or DNA is degraded (Clarke, 2014). Finally, there are financial advantages; while the initial setup for SNP genotyping may be costly, the cost per genotyping rapidly declines year-on-year (Ishige, 2023).

Due to these factors, parentage testing in numerous species has transferred from STRs to SNPs in the last two decades. As early as 2004, SNP parentage testing began on Holstein, Flechvieh, and Braunviesh cattle (Werner, 2004; Schutz, 2016). In 2012, the ISAG started SNP parentage comparison testing for cattle (Fernandez, 2013; Holl, 2017). Sheep parentage started the transition to SNP testing in 2014 (Heaton, 2014; Torereau, 2017). The Canadian bison industry began use of a SNP panel for parentage in 2020 (Yang, 2020). Cats and goats have also since switched to SNP-based parentage testing (de Groot, 2021; Talenti, 2021).

Despite the importance of horse pedigrees, the transition of equine parentage testing from STRs to SNPs has been slow. The first equine parentage SNP panel was carried out on Japanese thoroughbreds in 2010 using 53 SNPs, which found very low mutation rates compared to STRs (Hirota, 2010). These 53 SNPs were found to be as accurate as the 12 core ISAG STRs for horse parentage. In 2016, the KWPN Association in the Netherlands developed a warmblood SNP parentage test that also used the core markers from the 2010 SNP panel, supplementing it with additional SNPS (Arts, 2021). In 2017, a 670K SNP array was developed that tested parentage in 153 horses across 24 breeds (Schaefer, 2017). Also in 2017, 101 SNPs were tested across 729 horses representing 32 breeds (Holl, 2017). Further studies on thoroughbreds, draft horses, sport horses, Anglo-Arabians, and Arabian-Barbs were carried out between 2017-2024 (Flynn, 2021; Tozaki, 2021; Crichan, 2022; Ishaige, 2023; Aminou, 2024).

The success of these studies led the ISAG Equine Committee to perform pilot SNP parentage comparison tests, in addition to their core STR parentage markers, between 2017 and 2024. In total, the ISAG gave unanimous approval of 381 autosomal core SNPs plus 397 backup markers, with eight Y-chromosome and 76 X-chromosome SNPs at their 2023 Equine Committee Conference in Cape Town. These SNPs have been used for the ISAG equine *trial* SNP DNA Typing Comparison Tests since 2024.

However, there are potential problems with the ISAG approved SNPs. In the 2021 ISAG Equine SNP Comparison Test, concordance between laboratories was as low as 65.51% despite using the same set of SNPs. This was attributed to discrepancies between the different platforms used. This led to the Axiom MNEC670 array (now owned by ThermoFisher) becoming the recommended platform of all laboratories entering the 2024-2025 ISAG equine trial SNP comparison test (ISAG, 2024). However, given the upfront costs of array readers, this limits the number of laboratories when deciding to do parentage testing.

Moreover, as sequencing costs lower, laboratories, are increasingly adopting whole genome sequencing for comprehensive genetic testing, a trend already seen in other species. Making sure the SNPs selected for equine parentage are platform agnostic and valid for whole genome sequencing ensures further accuracy and allows laboratories to make the switch to whole genome sequencing as costs lower, providing greater choice for equine users. To date, the ISAG approved equine parentage SNPs have not been tested with whole genome sequencing.

Therefore, we performed the current study to analyze the core and backup ISAG SNPs by whole genome sequencing. Our results demonstrate that the use of platform-agnostic SNPs has transformative potential beyond parentage. They enable population structure analysis, breed determination, and ancestry, as well as performance traits and disease. It also allows for the integration of previously deposited historical genomic data, even when done on smaller, older SNP chips (Illumina EquineSNP50 and Illumina EquineSNP70), into the analysis. This allows for data from previous studies, encompassing thousands of horses, to still be utilized for research, while ensuring compatibility with the latest sequencing technologies.

## MATERIALS AND METHODS

### Sample Collection and Sequencing Datasets

To comprehensively evaluate the performance of different platforms to assay ISAG established parentage SNPs across multiple breeds, we used 833 equine samples representing 24 distinct breeds, taken from multiple sources. This dataset integrated both publicly available whole genome sequencing (WGS) data and SNP array genotypes, ensuring broad representation of genetic diversity and sequencing methodologies.

Public WGS were obtained from the National Center for Biotechnology Information and European Nucleotide Archive (ENA). These included 88 horses representing diverse breeds (ENA accession: PRJEB14779), sequenced on multiple Illumina platforms including HiSeq 2000, HiSeq 2500, HiSeq 3000, and NovaSeq 6000, providing essential platform diversity for validation. An additional 186 Thoroughbred VCF files generated using whole genome sequencing were obtained from the University of Kentucky Equine Genomics Center, of which 21 of these genomes are Japanese Thoroughbreds. The inclusion of WGS data from multiple sequencing centers was deliberate, ensuring sequencing platform and bioinformatic pipeline diversity to strengthen the broad validation of our findings.

Additional SNP array data from 364 public samples was used (https://figshare.com/s/a04031d1ab0fcebf7bee). This dataset comprised 172 horses genotyped on the GGP Equine 70K array (Illumina Inc.), representing 67 Turkoman, 39 Dareshuri, 35 Caspian, and 31 Kurdish horses (Ardestani, 2022). It also included an additional 192 samples genotyped from previous studies using the Axiom Equine Genotyping Array 670K (Affymetrix), representing 109 Iranian Asil, 9 Turkoman horses, 64 Kurdish, 5 Caspian, and 5 Dareshuri (Sadeghi et al., 2019; Yousefi-Mashouf et al., 2021). In total, 738 publicly deposited samples sequenced on 6 platforms were used from three data sources.

To complement the publicly available resources, we generated WGS for 195 horses. Blood and hair samples were collected using standard techniques, with blood drawn from the jugular vein and hair follicles collected according to established protocols. DNA isolation was performed using Sbeadex DNA Purification Kit (Biosearch Technologies) and QIAmp DNA extraction (Qiagen) according to the manufacturers’ instructions. Genomic DNA was sequenced using TruSeq DNA Nano (Illumina) library on the Illumina NovaSeq 6000 platform with 30x / 60 GB output at CeGaT GmbH (Germany). Additional sequencing was performed using the AVITI Cloudbreak Freestyle (Element Biosciences) library on the Element Biosciences AVITI System platform at Medlabs (Jordan). This achieved an average sequencing depth of 29X. Different laboratories, providers, libraries, and sequencing platforms were chosen to validate our platform agnostic approach. All sequencing data were processed on Amazon Web Services using the Illumina DRAGEN pipeline (https://aws.amazon.com/solutions/partners/illumina-dragen/) with alignment to the EquCab3.0 reference genome. Read mapping and variant calling were performed using the DRAGEN Pipeline with default parameters. This depth of coverage gave reliable variant calling across the genome, providing high-confidence genotypes for subsequent analyses.

### ISAG SNP Panel Evaluation

The ISAG approved 775 autosomal SNPs at their 2023 Equine Committee Conference in Cape Town, comprising 378 core SNPs and 397 backup SNPs. These positions were originally ascertained from commercial SNP arrays manufactured by Affymetrix and Illumina. While the ISAG panel also includes X and Y chromosome markers for quality control purposes, our analysis focused exclusively on autosomal SNPs for consistent evaluation across all samples regardless of sex.

### Bioinformatic Analysis

All genomic coordinates from downloaded datasets were standardized to the EquCab3.0 reference assembly using the UCSC LiftOver tool, ensuring consistent positional information across multiple data sources. Our internally generated whole genome sequences were processed using the Illumina DRAGEN Bio-IT Platform version 4.0.

Population structure and genetic relationships were assessed using the SNPRelate package in R, implementing principal component analysis (PCA) to visualize breed clustering and validate marker informativeness. Linkage disequilibrium patterns were calculated to ensure independent segregation of selected markers. Data visualization was performed using ggplot2 and RColorBrewer packages, generating figures for comprehensive representation of genetic relationships.

### Parentage testing with 1209 SNP Panel

VCF files were first filtered to retain only the 1,209 high-quality SNP positions using a custom filtering script. A perl script then performed pairwise comparisons between all 195 samples at these positions to identify potential parent-offspring relationships. The script applied the rule that if a parent is homozygous at a given position (either REF/REF or ALT/ALT), the offspring cannot be homozygous for the opposite allele. Specifically, if the parent genotype was REF/REF, the offspring could not be ALT/ALT, and vice versa. Following ISAG published guidelines, up to 5 mismatches were permitted at this stage to account for technical errors in sequencing or variant calling.

A Python script processed the candidate parent-offspring pairs from Stage 1 to assemble complete trios and apply stringent validation criteria. The script first confirmed that putative sires were male, and dams were female based on genetic sex determination. For each assembled trio, the script applied the ISAG rule that when both parents are homozygous for the same genotype at a position (both REF/REF or both ALT/ALT), the offspring cannot be heterozygous (REF/ALT). At this validation stage, 0-6 mismatches were permitted per ISAG recommendations, ensuring genetic consistency across all SNP positions.

## RESULTS

### SNP Classification and Quality Assessment

Stringent quality criteria were established to identify SNPs suitable for cross-platform parentage testing. VCF files from our 195 whole genome sequencing samples were first combined into a single VCF (gVCF) file and filtered for variants receiving a PASS filter using bcftools, indicating the variant caller’s confidence of 99.99% accuracy. This initial quality filtering produced minimal reduction, from approximately 5 million to 4.9 million variants, retaining only high-confidence calls. The PASS designation integrates read quality, base quality, mapping quality, and additional parameters used by variant caller that are specific to each genomic region.

These quality-filtered variants were then integrated with three additional datasets: 274 publicly available whole genome sequences and 364 SNP array samples (Figure 2). During this integration, only positions present in at least 10% of the samples were retained, yielding 28859 high-quality positions. This requirement meant that selected SNPs could be reliably detected regardless of platform or pipeline. For array data conversion to VCF format, duplicate positions representing multiallelic sites were resolved to single biallelic calls, as whole genome sequencing inherently represents each genomic position as a single record.

**Figure 1:**
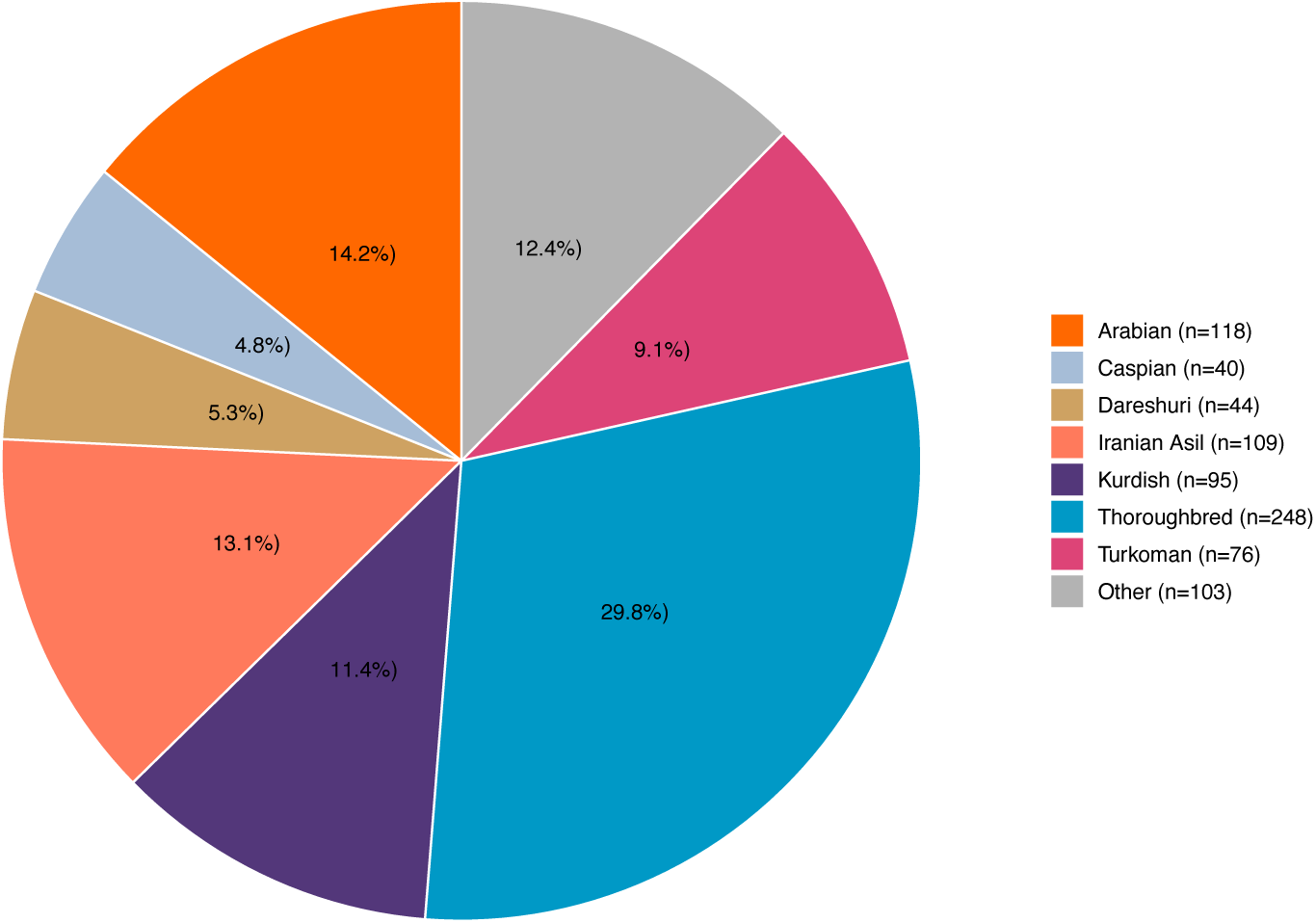
Distribution of 833 horse samples across different breeds.

**Figure 2:**
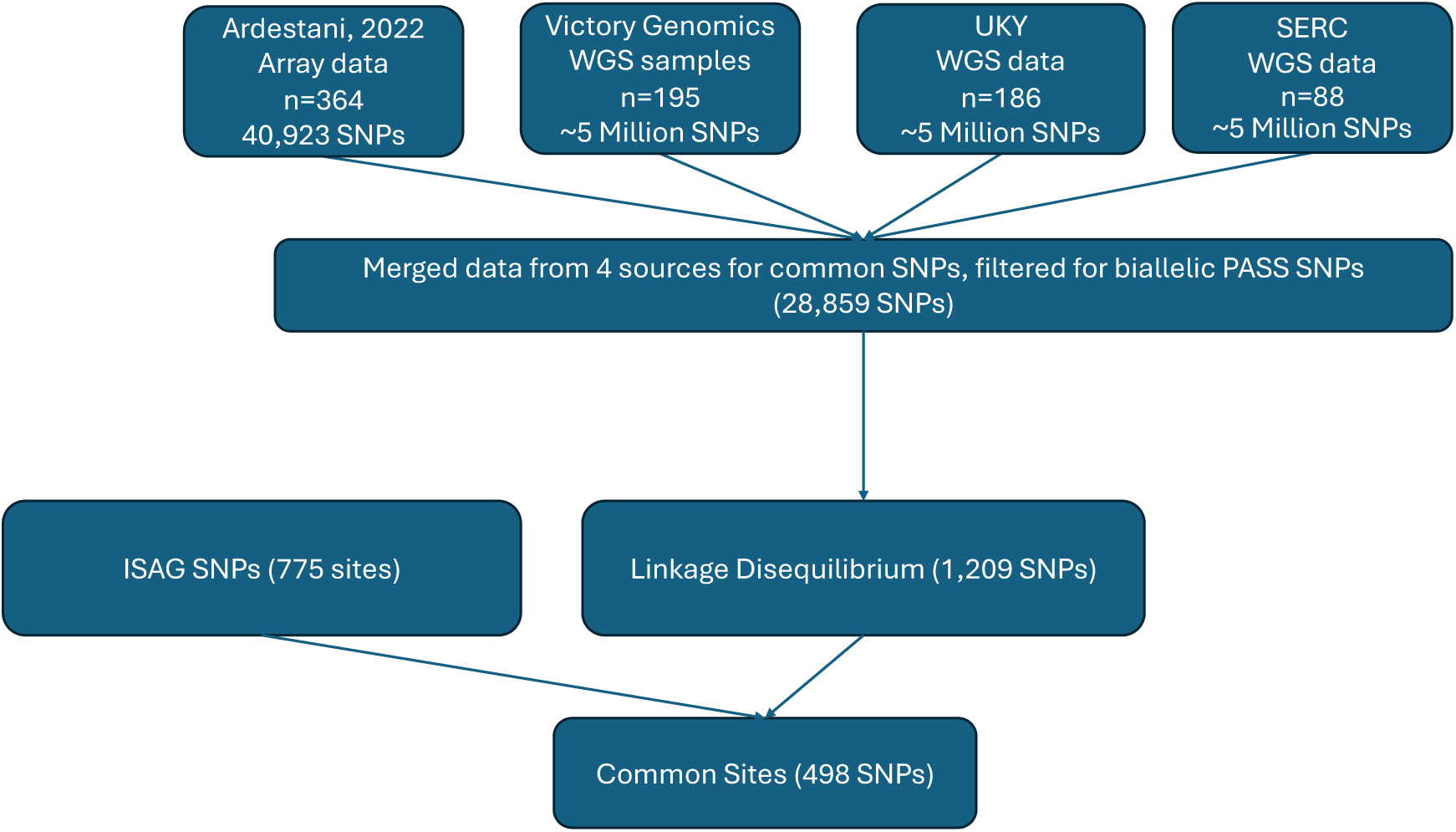
Flowchart of SNP filtering to establish a platform independent panel

The 28859 positions underwent linkage disequilibrium analysis for independent segregation of markers, which reduced the panel to 1,209 SNPs. This filtering step maximized the information content while eliminating redundant markers in high linkage. The final 1,209 SNPs demonstrated balanced distribution across all chromosomes, maintaining comprehensive genomic coverage for accurate parentage determination even in cases of regional technical failure.

### Development of Platform-Agnostic SNP Panel

Backward compatibility with existing infrastructure was prioritized throughout the selection process. Since our analysis incorporated data from both llumina Equine SNP70 and Axiom 670K arrays, backward compatibility was intrinsically validated. Each candidate SNP position underwent verification for reliable detection across all whole genome sequencing datasets, ensuring robust performance independent of sequencing depth or pipeline specifications.

Importantly, the final panel demonstrated balanced distribution for the 1,209 high quality selected positions across all chromosomes (Figure 3). This distribution gives comprehensive coverage while maintaining sufficient marker density for accurate parentage determination even in the presence of regional dropout.

**Figure 3:**
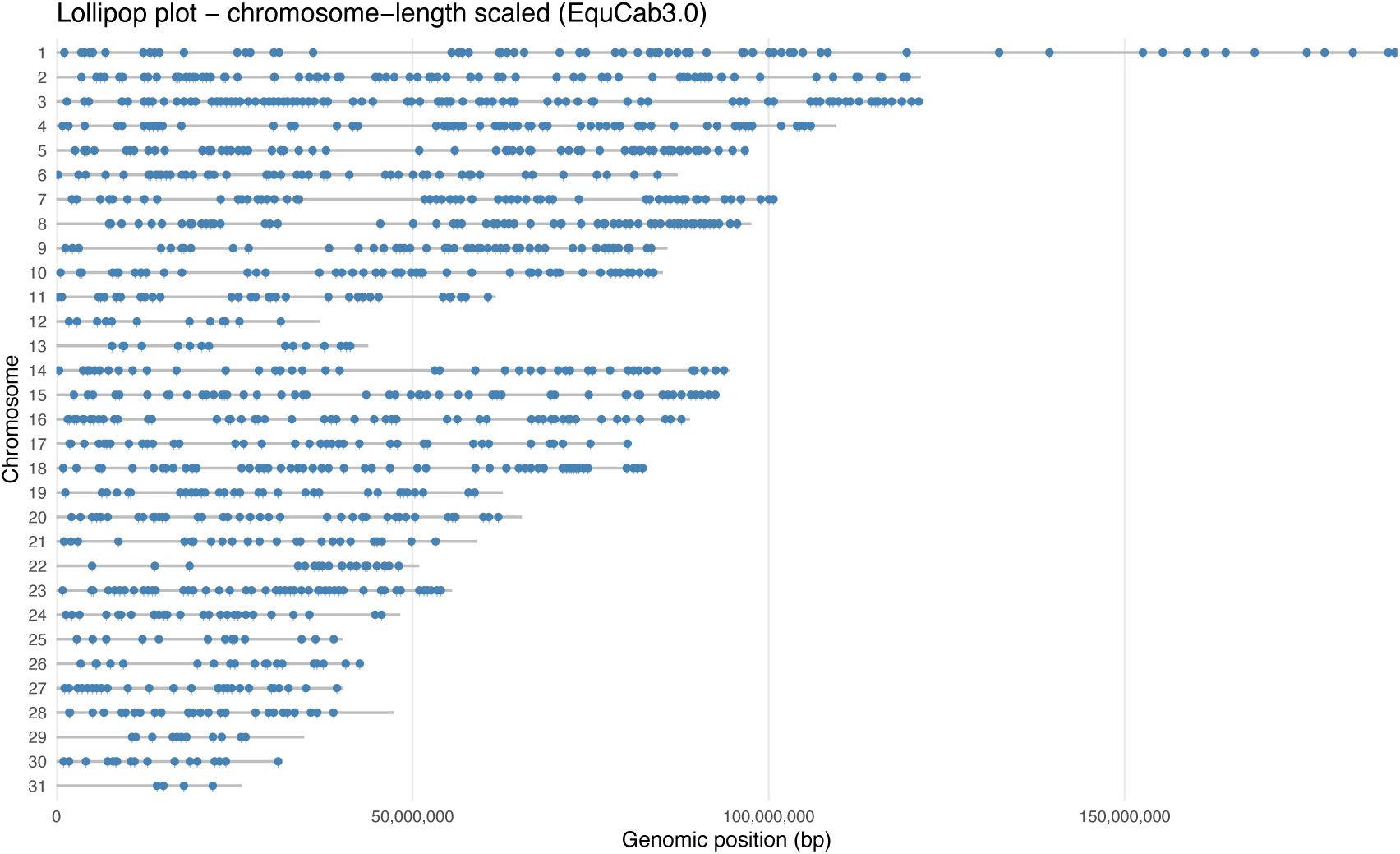
Chromosome distribution of proposed 1209 SNPs in this study

### ISAG SNP Performance in Whole Genome Sequencing

Analysis of WGS data for the ISAG-recommended parentage panel revealed substantial detection failures. Of the 775 autosomal ISAG-approved SNPs, 277 (35.7%) failed to meet quality thresholds across multiple WGS platforms. This failure rate remained consistent across independent datasets, with the same SNPs repeatedly showing inadequate quality scores or insufficient read depth. The distribution of failed SNPs showed no regional clustering, with problematic markers scattered across all equine chromosomes rather than concentrating in specific genomic regions (Figure 4). Breakdown by functional categories demonstrated that failures affected both core and backup SNP classifications equally, indicating the issue was not limited to supplementary markers.

**Figure 4:**
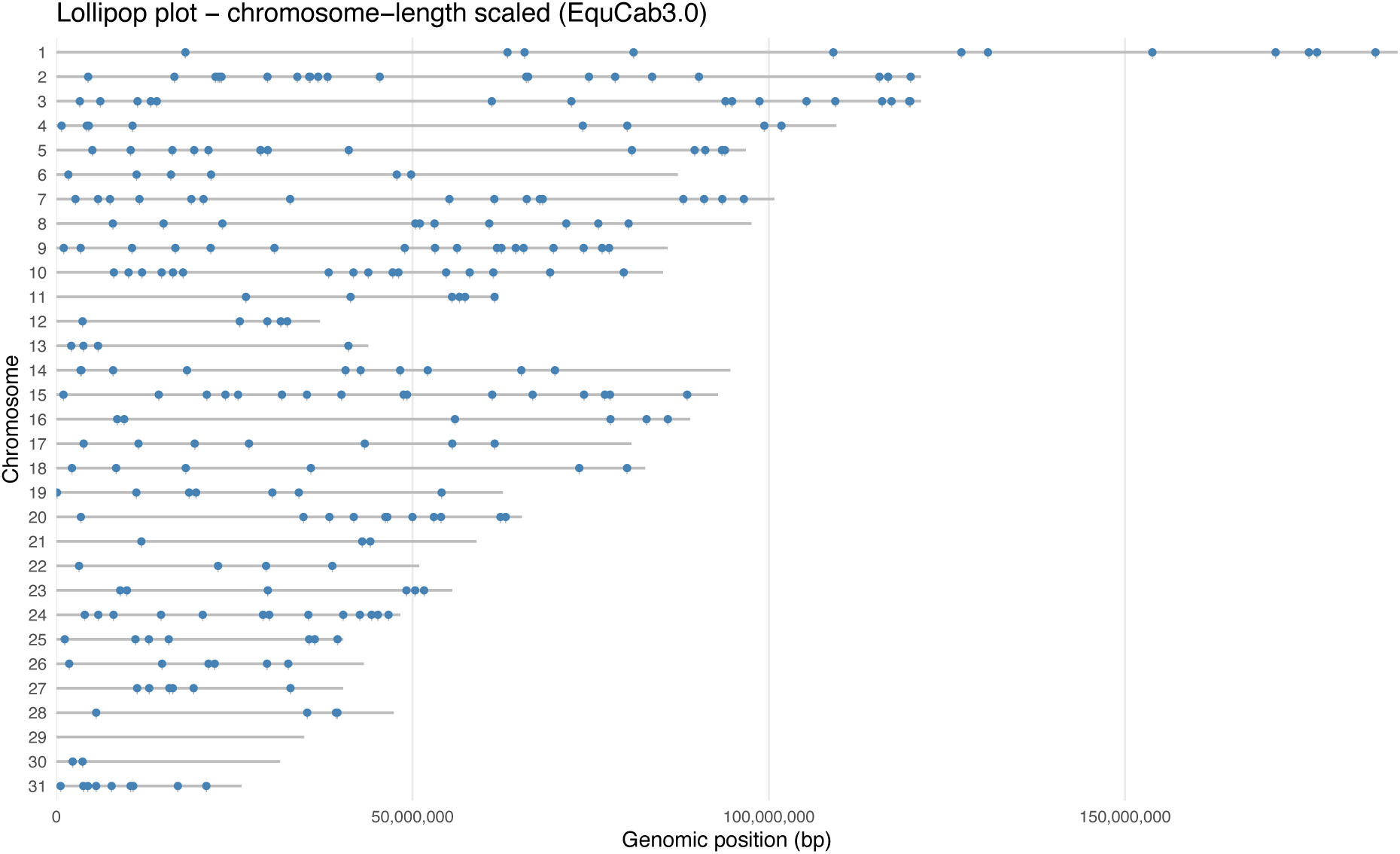
Distribution of ISAG SNPs not detectable by WGS from multiple laboratories

**Figure 5:**
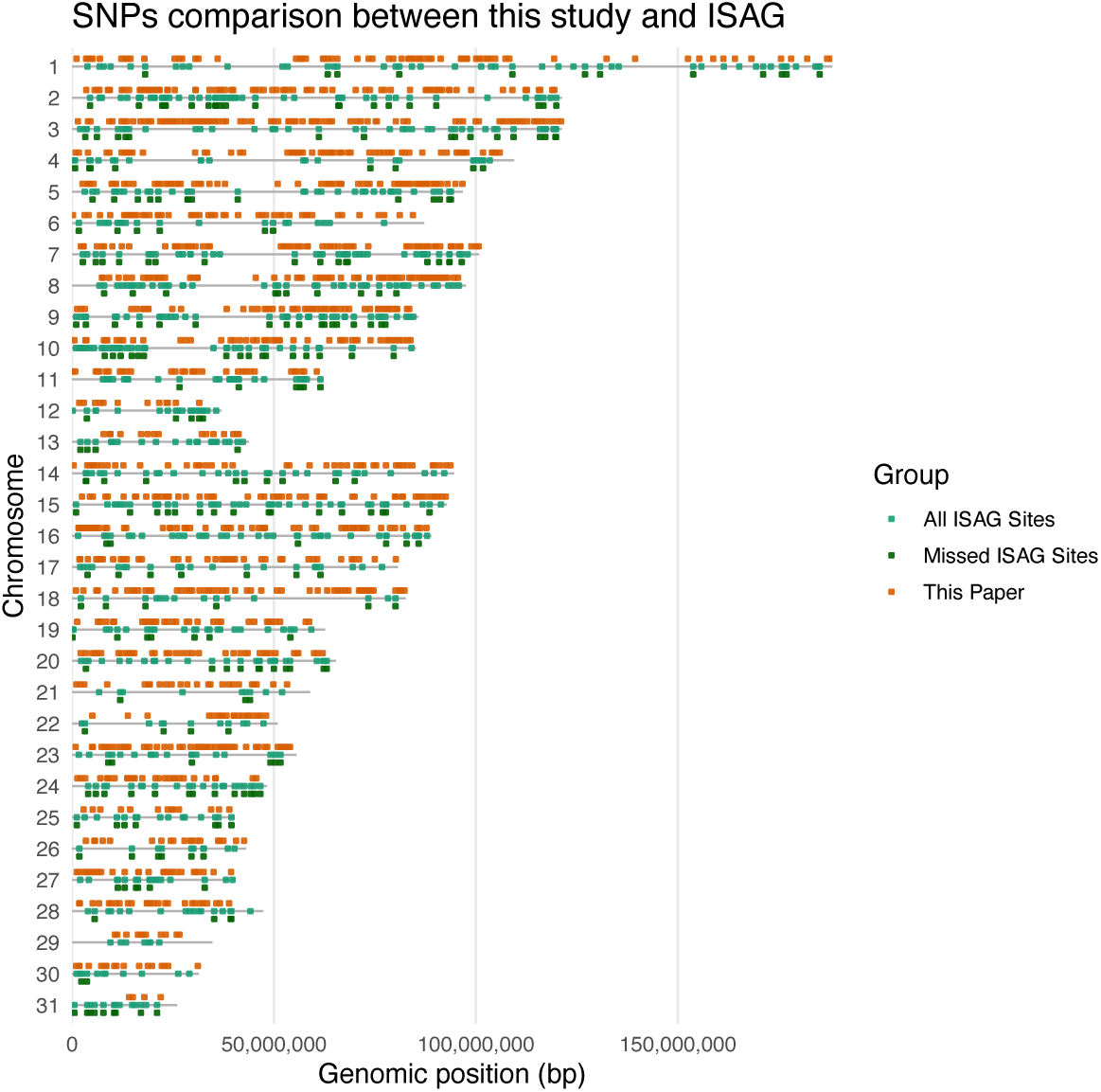
Distribution of all ISAG SNPs, ISAG SNPs not detectable by WGS, and the 1209 SNPs proposed by this paper

### Characterization of SNP Failures

Investigation into the root causes of SNP failures revealed a clear pattern of technical incompatibility between array-designed markers and WGS platforms. The primary failure mechanism involved poor short-read alignment in repetitive or structurally complex genomic regions. At these problematic sites, the DRAGEN quality scores indicated approximately 10% error rates in SNP calling, with read depths frequently falling to 1X coverage. While array hybridization successfully captures these regions through probe-based detection, short-read sequencing alignment consistently fails to generate reliable genotype calls at these positions.

Critical validation using publicly available VCF files from other research groups confirmed that these failures were not specific to our internal sequencing protocols or bioinformatics pipeline. The same 277 SNPs were absent or failed quality filters in WGS data generated by independent laboratories using different sequencing machines and analysis pipelines. This consistency across multiple data sources, sequencing platforms, and computational approaches demonstrated that the incompatibility represents a fundamental limitation of these specific SNP positions when interrogated through short-read whole genome sequencing methodologies.

### Comparative Performance Analysis

Direct comparison between the ISAG panel and our proposed 1,209 SNP panel demonstrated marked differences in platform compatibility. While 35.7% of ISAG autosomal SNPs failed quality thresholds in WGS applications, our platform-agnostic panel maintained consistent detection across all evaluated platforms. The 1,209 SNPs showed compatibility across the GGP 70K array, Axiom 670K array, and all WGS datasets tested, providing standardized performance regardless of genotyping technology employed.

Concordance testing performed across different sequencing platforms and multiple alignment pipelines confirmed the accuracy of our panel design. This comprehensive validation demonstrated that failures in the ISAG panel occur consistently across publicly available datasets processed through independent pipelines, while our proposed markers maintain quality scores suitable for analysis.

Importantly, 498 SNPs from the original ISAG panel worked across all platforms and overlapped with our panel (Figure 2), demonstrating that the ISAG selection process does not require complete revision but rather targeted replacement of problematic markers. Importantly, replacement of these markers does not require any redesign across either array or short-read sequencing platform, simply a reassignment of markers already backward compatible.

### Validating the 1209 SNPs for Parentage

To assess the practical application of our 1,209 SNP panel for parentage testing, we analyzed 195 whole genome sequencing samples to identify complete parent-offspring trios.

All 12 genetically determined trios (from 195 WGS samples) were cross-referenced against pedigree records and owner registration documents. The results demonstrated 100% concordance, with all trios correctly matching their registered pedigree information. This validates that the 1,209 SNP panel maintains the accuracy standards required for parentage verification in equine populations when applied to whole genome sequencing data.

### Population Structure Analysis

With STR data, little information about the horses analyzed can be inferred due to the relatively small amount of data generated. Effectively, STR markers can generally define parentage and not much more. However, we hypothesized that we could infer additional information from SNP panels.

We began with an analysis of population structure. We first examined the 498 ISAG SNPs that overlapped with the 1,209 SNPs we identified and are compatible across all platforms in all 833 samples representing 24 breeds, using Principal Component Analysis (PCA), a visualization method for data similarity. Therefore, points adjacent to one another in PCA have datasets that are more similar than those points more distant. In the context of equine genomes, PCA enables detection of close relationships between horse genotypes from more distant ones, each point representing the cumulative genotype of one horse. Using PCA, there was a general separation of Thoroughbreds from Arabians, although there was substantial overlap across multiple horse breeds when examining these 498 ISAG SNPs (Figure 6).

**Figure 6:**
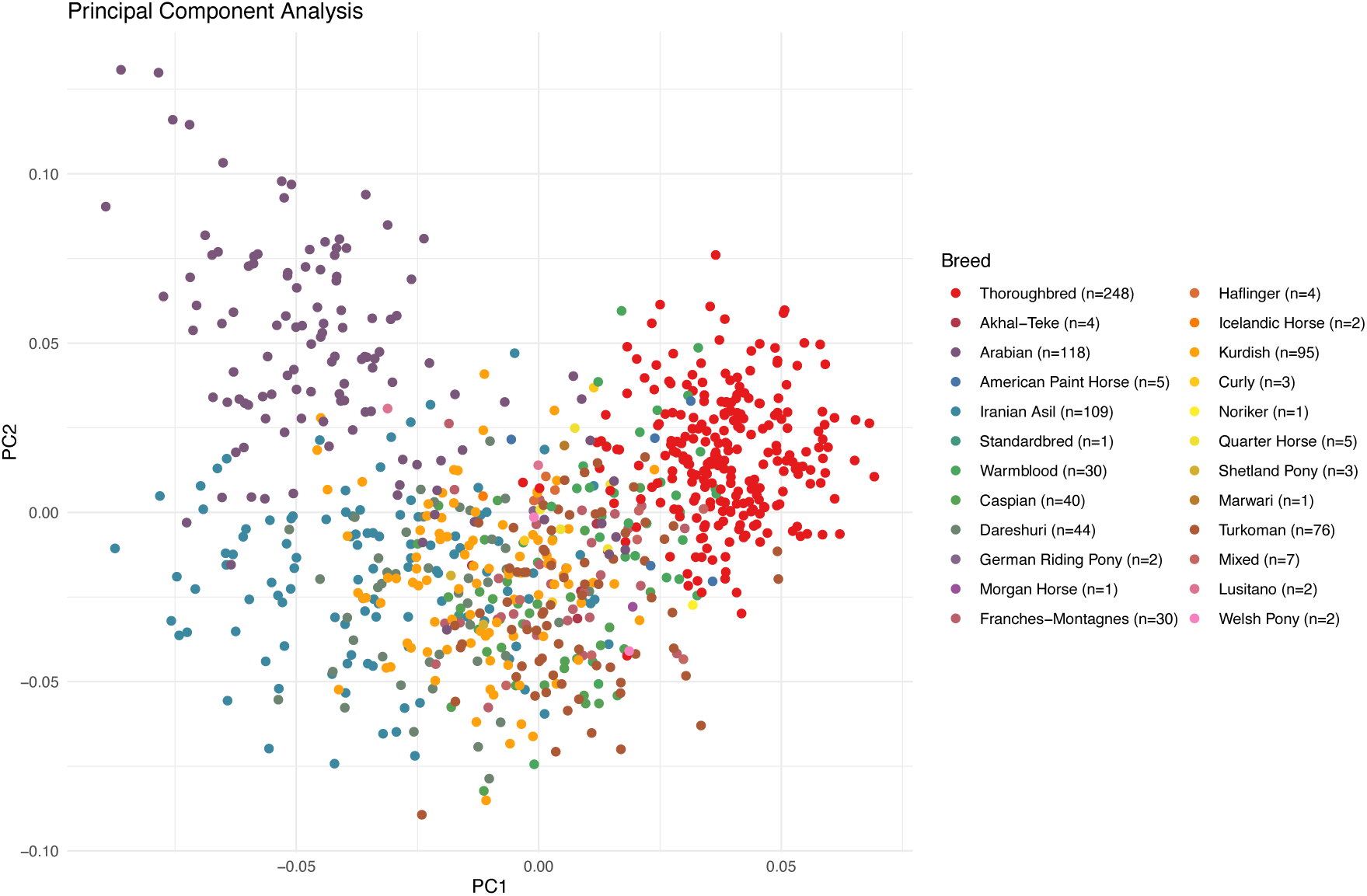
Population structure using ISAG SNPs available across platforms

In contrast, when our proposed 1209 platform-agnostic SNPs were used, the Arabian horses demonstrated clear separation to Iranian Asil and other native Middle Eastern breeds (Figure 7). Thoroughbreds formed a very distinct cluster, while Warmblood breeds occupied an intermediate genetic space as expected from their known breeding history. Breed assignment accuracy was validated in 195 WGS samples through phenotype information, breed registry documentation, and pedigree records supplied by owners. The analysis successfully identified registered “pure” Arabians that showed evidence of Thoroughbred influence, demonstrating the panel’s utility beyond simple parentage verification for detecting admixture and verifying breed purity claims.

**Figure 7:**
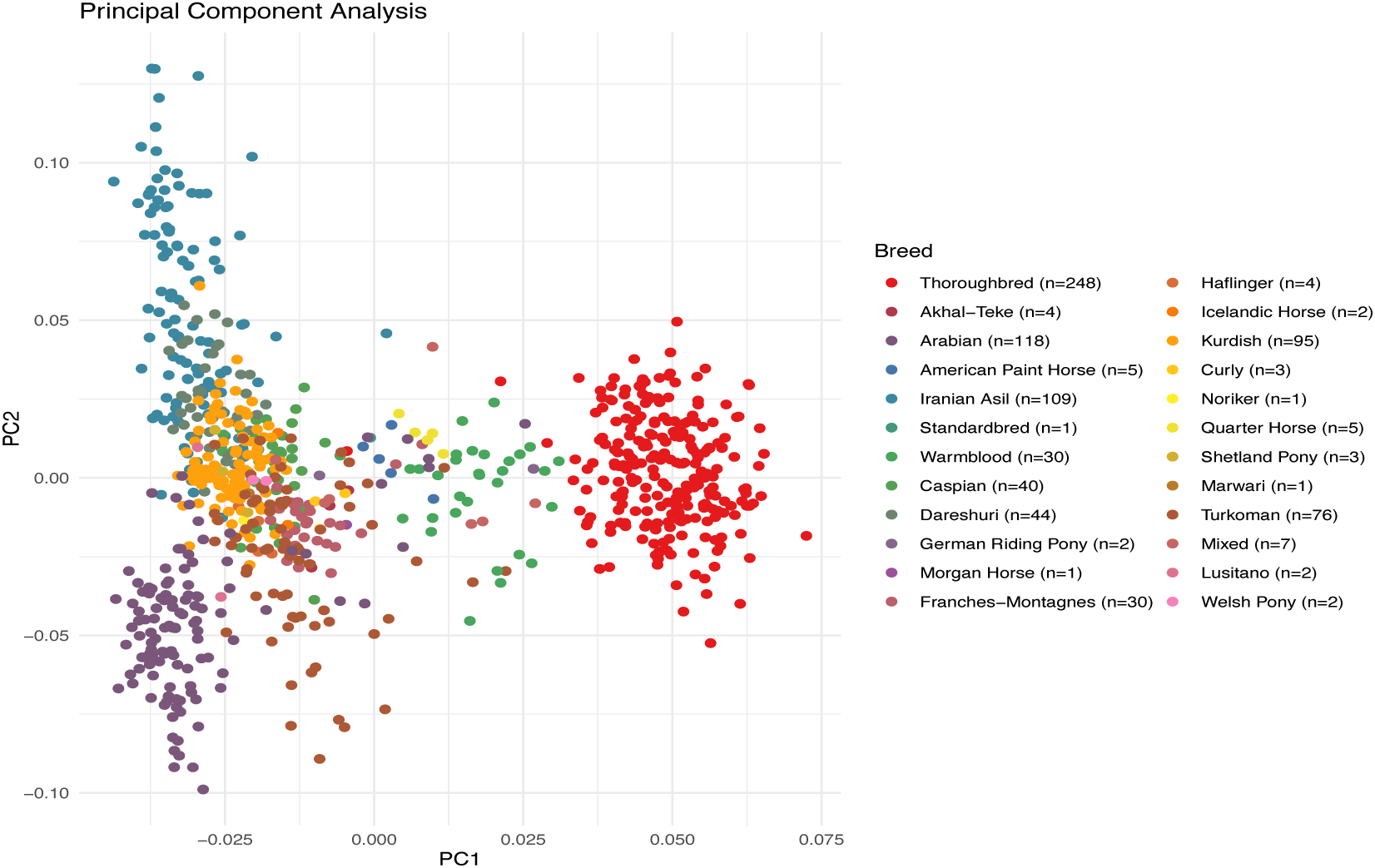
Population structure using proposed 1209 cross-platform SNPs

Finally, we examined the 28,000 SNPs that are common across all the microarray data and WGS, representing the largest genotype dataset that is common across all 833 samples (Figure 8). PCA analysis demonstrated even greater resolution across breeds illustrating the value of extended SNP data, such as from WGS, for resolving relationships across breeds.

**Figure 8:**
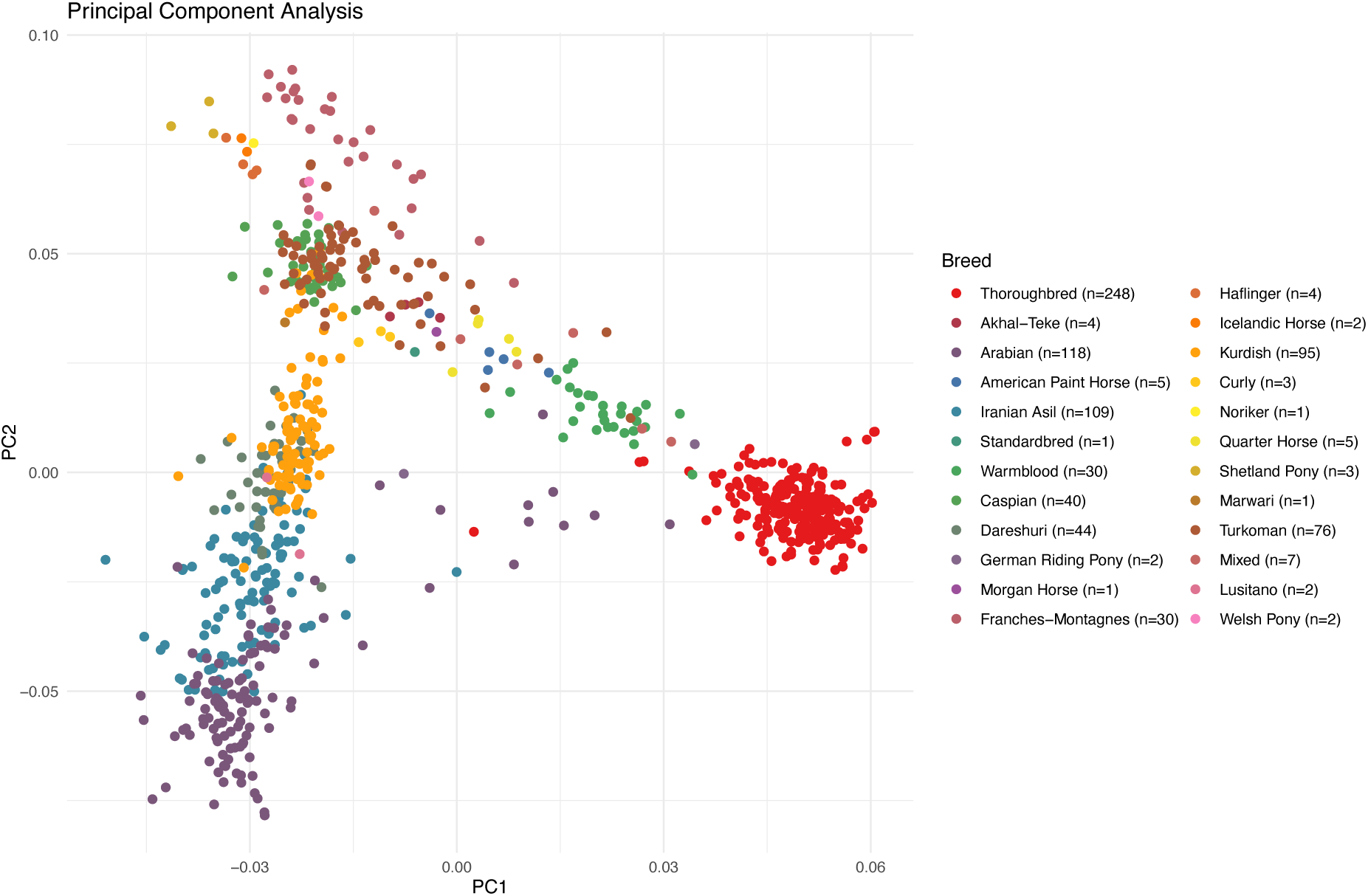
28K filtered SNPs in common between WGS and existing equine SNP arrays

## DISCUSSION

In this study, we demonstrate that 277 (35.74%) of the 775 ISAG-recommended SNPs for equine parentage testing cannot be accurately called by short-read, whole genome sequencing despite high-depth and high-coverage. This finding is significant as short-read next generation sequencing such as WGS is becoming increasingly widespread. Indeed, the cost of WGS is plummeting enabling this technology to be applied to multiple animal species. As such, millions of SNPs and other sequence variants can be analyzed to better understand the genetic architecture of animal traits which make these animals so valuable.

Our results were strengthened by the use of multiple, independent, public WGS datasets sequenced on different platforms as well as our own WGS data. In addition, there was consistent failure across different bioinformatic pipelines.

Therefore, this platform incompatibility cannot be attributed to specific sequencing technologies or bioinformatics pipelines. In short, the proposed ISAG equine parentage SNPs designed for array or chip platforms encounter systematic issues in repetitive or complex regions of the genome and are not suitable for short-read WGS, the predominant sequencing platform.

To address this issue, we selected 1209 high quality SNPs. These 1209 SNPs were found in 24 horse breeds and distributed across all chromosomes (not including X/Y). Moreover, all these SNPs have backward compatibility being available on all existing equine SNP arrays and chips.

As equine SNP genotyping arrays has progressed through several generations (Illumina EquineSNP50; Illumina Equine SNP70; Axiom MNEc2M; Axiom MNEc670K), we confirmed that the 1209 SNPs have backward compatibility being available on all existing equine SNP arrays and chips, as well as forward compatibility with WGS. One of the remits in designing new equine SNP arrays to supercede previous generations has been backward compatibility so that historical datasets, regardless of platform, could still be used (Schaefter, 2020). However, the currently proposed ISAG parentage SNPs do not allow for *forward* compatibility in that 35% of them cannot be called on WGS platforms that are increasing in popularity. Therefore, our proposed 1209 SNPs allows forward compatibility with WGS platforms. The selection of these high-quality SNPs, distributed across all chromosomes, and tested across many breeds, ensures accurate horse parentage calling using SNPs regardless of platform, bioinformatics pipeline or laboratory. It also gives assay providers and horse owners a choice of technology platform.

Importantly, however, the efforts of the ISAG over the last 15 years in transitioning to SNPs for equine parentage is welcomed and should not be downplayed. Our aim is not to dismiss but to augment this work, which has been a crucial step to more accurate, multiplexed, faster, and cheaper parentage confirmation. On this point, we found that 498 ISAG selected parentage SNPs can still be used with WGS, as they are encompassed within the 1209 high quality SNPs outlined in this study.

Moreover, as part of our proposed platform agnostic approach, all 1209 SNPs in this study are found on the Thermofisher (Axiom) 670K array preferred by ISAG. This has the advantage that no new chip or array development is needed. Instead, the 1209 platform agnostic SNPs could be recommended by ISAG and be used on the current array or any other.

There are indications, however, where the richer dataset offered by WGS has advantages. Population structure analysis has practical applications in breed verification for registration, detection of crossbreeding, identifying horses by use, and assessment of genetic diversity within breeding programs. Our PCA analysis showed that with the 498 ISAG SNPs that can be assayed regardless of platform, breed separation was minimal with overlapping clusters providing little discriminatory power between populations. This changed dramatically when using our 1,209 platform-agnostic SNPs, where distinct breed clusters emerged with clear separation between populations. This enhanced resolution for breed identification and population stratification is obviously expected, as more markers means more resolution. However, the improvement was noticeable and could greatly improve our understanding of population structure.

While laboratories may use larger SNP arrays containing thousands or tens of thousands of markers for enhanced population analysis, this approach faces economic constraints. Each additional SNP included on an array increases reagent costs, data processing requirements, driving up the per-sample testing price. The 1,209 platform-agnostic SNPs have sufficient markers for basic population analysis while remaining economically viable for routine testing.

For population analysis and other genomic discovery, WGS is a superior platform because so much more data is generated. As equine science gradually transitions to WGS, having a platform-agnostic standardized SNP panel becomes even more urgent. Not to do so would mean that researchers using WGS would not be able to confirm parentage using the current equine SNPs.

## CONCLUSION

In genomics, across all species, there is a gradual move towards whole genome sequencing. This is facilitated by the year-on-year cost reductions of WGS, due to a multitude of innovations and new sequencing platforms and providers. WGS also means more data availability for research. This aids the discovery of variants for genetic disease, many traits, better inbreeding calculations, the influence of mtDNA, conservation, and more. Advances in AI and Machine Learning are greatly enabled by the rich data from WGS for further discovery. It is almost certain that in coming years WGS will become more utilized by research and commercial groups. Should the ISAG require equine parentage SNPs that cannot be called using WGS, then the ISAG faces the prospect of redoing their current SNP parentage panel in the near future.

To emphasize, the work of the ISAG in moving towards SNP parentage testing is welcome, as is the contributions of various equine research groups to this process. However, this is the first time these SNPs have been tested on WGS resulting in 277 not being called on this platform. This study defines 1209 high-quality SNPs, which also encompass 498 of the proposed ISAG SNPs, forming a comprehensive, platform-agnostic equine parentage panel that does not require a redesign of the current equine 670K SNP array while being simultaneously suitable for WGS.

## PUBLIC DATA SOURCES

186 WGS samples from the University of Kentucky: https://equinegenomics.uky.edu/thoroughbredDiversity.html

364 SNPs deposited by Ardestani (2022) https://figshare.com/s/a04031d1ab0fcebf7bee

88 horses representing diverse breeds (ENA accession: PRJEB14779)

For reasons of client confidentiality, we may not be able to provide the whole genome but can supply SNPs on reasonable request. Please contact the corresponding author.

